# Cell-resolved pharmacodynamic profiling reveals tissue- and cell-population-specific activity of antisense oligonucleotide chemical conjugates *in vivo*

**DOI:** 10.64898/2026.10.30.762893

**Authors:** Shuzhi Cheng, Jennifer Frommer, Ayad Eddaoudi, Panayiota Constandinou, Matthew Wood, Francesco Muntoni, Haiyan Zhou

**Affiliations:** Genetics and Genomic Medicine Research and Teaching Department, Great Ormond Street Institute of Child Health, University College London, London WC1N 1EH, UK; Department of Paediatrics, University of Oxford, Oxford OX3 9DU, UK; Flow Cytometry Core Facility, UCL Great Ormond Street Institute of Child Health, University College London, Zayed Centre for Research into Rare Disease in Children, London WC1N 1DZ, UK; Neurodegenerative Disease Department, UCL Queen Square Institute of Neurology, London WC1N 3BG; Developmental Neurosciences Research and Teaching Department, Great Ormond Street Institute of Child Health, University College London, 30 Guilford Street, London WC1N 1EH, UK; NIHR Great Ormond Street Hospital Biomedical Research Centre, London WC1N 1EH, UK

**Author notes:** Correspondence to: Haiyan Zhou Great Ormond Street Institute of Child Health, University College London, 30 Guilford Street, London, WC1N 1EH, United Kingdom. These authors contributed equally (Shuzhi Cheng, Jennifer Frommer).

**Keywords:** antisense oligonucleotide, gapmer ASO, ASO conjugate, *Malat1*, target engagement, biodistribution, PDGFRα, eGFP, FACS

## Abstract

Chemical conjugation can modulate antisense oligonucleotide (ASO) pharmacology and improve organ-specific delivery, yet whole-tissue measurements provide limited information on cell-specific activity. Here, we used *metastasis-associated lung adenocarcinoma transcript 1* (*Malat1*) knockdown as a pharmacodynamic readout to compare whole-tissue and cell-resolved activity of chemically conjugated ASOs in *platelet-derived growth factor receptor alpha-enhanced green fluorescent protein* (PDGFRα-eGFP) mouse model to assess their *in vivo* uptake in PDGFRα-expressing (GFP^+^) cells. This reporter mouse model uses PDGFRα as an exemplar cellular marker and provides an accessible tool for evaluating ASO conjugate activity within mesenchymal stromal and progenitor populations, mainly tissue fibroblasts. Following subcutaneous administration, *Malat1* was quantified in whole tissues and cell-sorted PDGFRα^+^/GFP⁺ or PDGFRα^-^/GFP⁻ populations. ASO-mediated *Malat1* reduction varied by tissue and conjugate chemistry, with palmitate conjugation producing the broadest extrahepatic activity and pronounced whole-tissue activity in heart and skeletal muscle. Cell-resolved analysis further revealed preferential palmitate-ASO activity within PDGFRα⁺ skeletal-muscle populations. Preliminary safety assessment also showed conjugate-dependent differences, with body-weight loss and ALT elevation in tocopherol-ASO-treated male mice and spleen enlargement in both sexes. These findings present an *in vivo* modelling system for evaluating cell-specific ASO activity and demonstrate the advantages of cell-resolved assessment over whole-tissue measurements.

## INTRODUCTION

Antisense oligonucleotides (ASOs) have emerged as a leading therapeutic approach for genetic diseases, enabling direct intervention at the RNA level and disease mechanisms that may not be accessible to conventional drug treatment.^1–4^ Specifically, gapmer ASOs contain a central DNA region flanked by affinity-enhancing chemically modified nucleotides and typically act through RNase H1-mediated cleavage of the target RNA after Watson-Crick hybridisation.^1, 2^ Introducing phosphorothioate backbones and sugar modifications such as locked nucleic acid, constrained ethyl, and 2’-*O*-methoxyethyl chemistries increases nuclease resistance, plasma protein binding while ensuring tissue exposure, enabling many ASOs to be administered without a delivery vehicle.^1, 2^

Despite these advances, targeted delivery remains a central limitation for broadening ASO therapeutic applications, particularly outside tissues that are naturally permissive to systemically administered oligonucleotides.^5–7^ Systemically administered ASOs preferentially accumulate in the liver and kidney, whereas productive delivery to tissues such as heart, skeletal muscle and lung remains challenging, particularly within specific cellular populations.^5, 8–11^ Despite successful clinical applications of intrathecal ASO delivery, systemic delivery to the central nervous system remains limited by the blood–brain barrier.^5^ ASO pharmacology is shaped by barriers that include vascular access, blood brain barrier, extracellular distribution and its potential alteration by pathological tissue remodelling (e.g., fibrosis), protein binding, cell-surface interaction, endocytosis, endosomal escape and access to the nuclear or cytoplasmic target RNA.^5, 6, 12–15^ Consequently, total tissue exposure, histological accumulation and productive RNA knockdown can diverge. Recent tissue pharmacokinetic studies further show that ASO exposure and knockdown vary by tissue, route of administration and conjugation chemistry, underscoring the need to interpret biodistribution together with functional target engagement.^6, 7^

Chemical conjugation is one of the most active strategies for modifying ASO distribution and potency. Ligands such as carbohydrates, lipids, peptides, antibodies and other biomolecules can alter plasma residence, allow receptor-mediated uptake, tissue penetration and intracellular trafficking.^5, 16–18^ GalNAc conjugation has transformed hepatocyte-targeted delivery, whereas extrahepatic targeting remains challenging and often depends on ligand, linker and tissue biology. ^5, 16–18^ Hydrophobic conjugates, including fatty-acid and tocopherol-based designs, can increase association with serum albumin or lipoproteins and modify tissue exposure, but increased accumulation does not always translate into proportional intracellular target engagement.^19–22^ For example, palmitic acid-conjugated *Metastasis-associated lung adenocarcinoma transcript 1 (Malat1)* ASOs alter plasma and tissue distribution in mice, which activity is shaped by endothelial transport and protein binding rather than by simple bulk accumulation alone.^19^. *Malat1* is a highly abundant, ubiquitously expressed nuclear RNA that is readily susceptible to RNase H-dependent gapmer-mediated knockdown, making it a widely used pharmacodynamic readout of ASO activity across tissues and cell types.^7, 23^

Importantly, a key limitation of whole-tissue homogenates is their inability to resolve cell-specific ASO activity. While informative at the organ level, it can obscure activity within specific cell populations, particularly in complex tissues where mesenchymal stromal, endothelial, immune, epithelial and parenchymal cells contribute differentially to bulk RNA measurements. The combination of cell-resolving technologies with ASO treatment has been used to assess *in vivo* cell-specific ASO delivery and targeting. For example, flow cytometry and single-cell RNA sequencing after intratracheal LNA gapmer delivery showed robust and durable gene silencing in lung fibroblasts and substantial activity in endothelial cells, establishing cell-resolved pharmacodynamic analysis as a useful approach for ASO delivery studies.^24^

Reporter-based models offer a complementary approach by enabling the isolation of genetically defined cellular compartments using a common fluorescence-based sorting strategy across multiple tissues. Although the abundance and recovery of reporter-positive cells vary between organs, this approach can reduce reliance on tissue-specific antibody panels and facilitate systematic comparison of ASO pharmacodynamic activity within defined cellular populations. Such analyses may reveal conjugate-dependent responses that are not apparent from conventional whole-tissue measurements, providing valuable insights to guide the development of extrahepatic ASO delivery strategies. Platelet-derived growth factor receptor alpha (PDGFRα) is a cell surface marker and functional regulator used to identify and isolate mesenchymal stromal and progenitor cells that play critical roles in tissue development, homeostasis, and fibrosis, including multi-potent or lineage-restricted progenitors capable of differentiating into various cell types, such as fibroblasts, myofibroblasts, and adipocytes.^25, 26^

Tissue fibroblasts and related mesenchymal stromal populations are increasingly recognised as therapeutically relevant cell types in a broad range of genetic and acquired disorders characterised by extracellular matrix dysregulation, tissue remodelling and fibrosis. In fibrotic disorders of the lung, liver, kidney and heart, including acquired conditions such as idiopathic pulmonary fibrosis, fibroblasts and related stromal populations participate in extracellular matrix deposition and tissue remodelling, making them attractive targets for anti-fibrotic therapies.^27^ In skeletal muscle, fibro-adipogenic progenitors (FAPs) have emerged as key regulators of regeneration, inflammation and fibrosis, and their persistent activation contributes to pathological fibro-fatty remodelling in muscular dystrophies such as Duchenne muscular dystrophy.^28–31^ In collagen VI-related muscular dystrophy, a group of autosomally recessive or dominant conditions due to mutations in either COL6A1, COL6A2 or COL6A3, and characterized by variable clinical severity, from infantile onset to onset in adolescent or adult life, fibroblast-lineage cells are a major source of collagen VI in skeletal muscle, providing a distinct rationale for targeting stromal populations.^32^ Importantly, PDGFRα expression does not define a single pure fibroblast subtype; rather, it labels tissue-context-dependent mesenchymal and fibroblast-like compartments.^30, 33^ Consequently, strategies capable of selectively delivering ASO therapeutics to PDGFRα-associated stromal populations could have broad translational relevance across multiple disease settings.

The present study was not designed to evaluate a disease-specific therapeutic target. Rather, we sought to establish a pharmacodynamic framework that could be used to assess future ASO delivery approaches directed towards fibroblast-enriched stromal populations. Using *Malat1* as the RNA target and functional pharmacodynamic readout, we compared ASOs bearing different chemical conjugations following subcutaneous administration in the PDGFRα-eGFP mouse model. The reporter system enabled cell-resolved assessment of productive ASO activity within PDGFRα-associated stromal populations *in vivo*. We analysed *Malat1* knockdown in whole tissues and in fluorescence-activated cell sorting (FACS)-isolated GFP^+^ and GFP^-^ populations from selected organs. By comparing bulk-tissue and reporter-defined cell populations for palmitate-, lactose- and tocopherol-conjugated ASOs, we aimed to determine whether different conjugate architectures redirect ASO activity at the tissue and cell-population levels *in vivo*, and whether whole-tissue pharmacodynamic readouts reflect target engagement within PDGFRα-associated reporter-defined cell populations.

## RESULTS

### Design and synthesis of distinct ASO conjugate architectures

To compare distinct conjugate architectures, all conjugates were based on the same RNase H-recruiting 2′-*O*-methoxyethyl (2′-MOE)/DNA gapmer containing a fully phosphorothioate (PS) backbone. The common ASO scaffold was functionalised with three chemically distinct biomolecules representing different ligand classes: the fatty acid palmitate, the lipid-soluble vitamin tocopherol (vitamin E), and the disaccharide lactose (Figure 1A). Palmitate and tocopherol were selected as representative lipophilic conjugates with established effects on ASO biodistribution, whereas lactose was included as an exploratory polar carbohydrate ligand. In contrast to GalNAc, which has become the benchmark carbohydrate conjugate for hepatocyte targeting, lactose has received little attention as a direct ASO conjugate.

**Figure 1.**
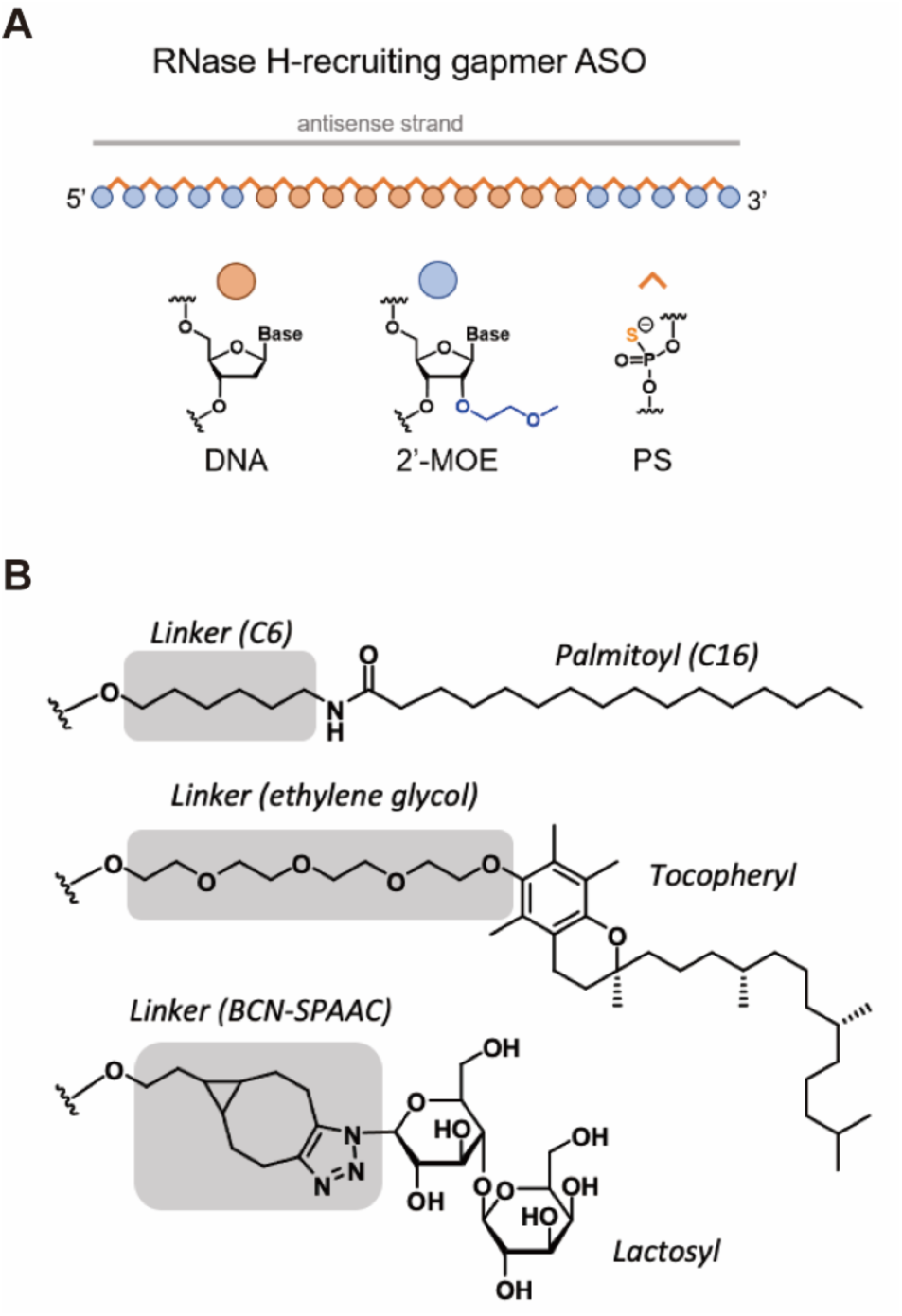
Chemical structures and conjugation strategies of the ASO evaluated in this study. **(A)** All conjugates were based on the same fully phosphorothioate (PS) 5–10–5 2′-O-methoxyethyl (2′-MOE)/DNA gapmer and differed in the attached biomolecule and linker architecture. **(B)** Palmitoyl and tocopheryl moieties were incorporated at the 5′ end through C6 and ethylene glycol linkers, respectively. Lactosyl-ASO was generated by post-synthetic copper-free strain-promoted azide– alkyne cycloaddition (SPAAC) between a 5′-BCN-modified ASO and azide-functionalised lactose, resulting in a triazole-containing BCN-derived cycloadduct. BCN, bicyclo[6.1.0]nonyne.

The three ASO conjugates differed not only in ligand identity but also in linker architecture and conjugation strategy (Figure 1A). Palmitate and tocopherol were introduced directly during solid-phase oligonucleotide synthesis using commercially available phosphoramidites, resulting in conjugates containing a C6 alkyl or ethylene glycol linker, respectively. In contrast, lactose required a post-synthetic conjugation strategy because no suitable lactose phosphoramidite was available. A bicyclo[6.1.0]nonyne (BCN) phosphoramidite was therefore incorporated at the 5′ end during oligonucleotide synthesis and subsequently reacted with azide-functionalised lactose by copper-free strain-promoted azide–alkyne cycloaddition (SPAAC), generating a stable triazole-containing BCN-derived cycloadduct between the ASO and lactose (Figure 1B).

Consequently, the three compounds represent distinct conjugate architectures in which ligand identity and linker composition vary simultaneously. While palmitate and tocopherol contain comparatively compact C6 and ethylene glycol linkers, respectively, SPAAC introduces a bulkier BCN-derived cycloadduct into the lactose conjugate. The present study was not designed to distinguish the individual contributions of ligand identity and linker architecture; the biological effects observed for each compound therefore reflect the properties of the complete conjugate rather than those of the attached ligand alone.

### FACS-based isolation of GFP⁺ and GFP⁻ cell populations from organs isolated from PDGFRα-eGFP mice

To assess tissue- and cell-population-specific ASO target engagement *in vivo*, PDGFRα-eGFP mice were treated by a single subcutaneous injection of ASO and conjugates. Tissues, including liver, kidney, spleen, heart, lung, skeletal muscle, sciatic nerve, spinal cord, skin and brain, were collected 5.5 days after for parallel whole-tissue and FACS-based analyses (Figure 2A). A subset of tissues was processed into single-cell suspensions for flow cytometry and sorting, whereas a broader tissue panel was collected for whole-tissue RNA analysis. Sorted GFP⁺ (i.e. PDGFRα^+^) and GFP⁻ (i.e. PDGFRα^-^) populations, together with whole-tissue samples, were subsequently analysed by RT-qPCR for *Malat1* expression as a functional readout of ASO target engagement. Body weight monitoring and plasma biochemical analysis were included as exploratory safety assessments.

**Figure 2.**
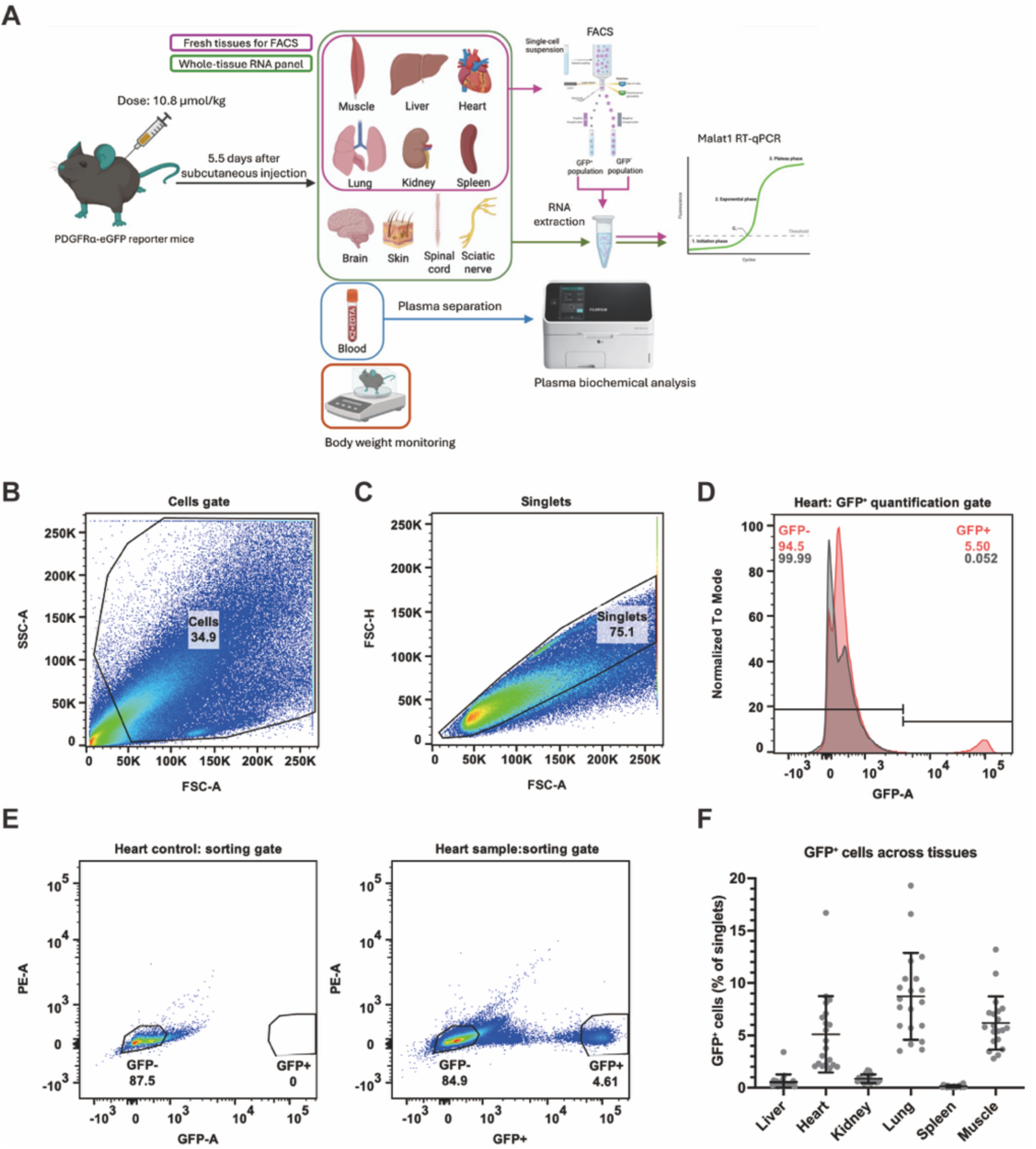
Experimental workflow and FACS-based separation of GFP⁺ and GFP⁻ tissue-derived populations from PDGFRα-eGFP mice. **(A)** Schematic overview of the in vivo experimental workflow. PDGFRα-eGFP reporter mice received subcutaneous ASO administration at 10.8 μmol/kg. Tissues were collected 5.5 days after injection for whole-tissue RNA analysis and/or preparation of single-cell suspensions for FACS. GFP⁺ and GFP⁻ populations were isolated from selected tissues, followed by RNA extraction and Malat1 RT-qPCR analysis. Body weight monitoring and plasma biochemical analysis were also performed. **(B–C)** Representative gating plots generated by post-acquisition analysis of heart-derived FCS data using FlowJo. Cells were gated using FSC-A and SSC-A (B), followed by singlet discrimination using FSC-A and FSC-H (C). **(D)** Representative histogram-based GFP gate used to quantify GFP⁺ cell frequency as a percentage of singlet events. The red histogram represents the PDGFRα-eGFP heart sample and the grey histogram represents the matched control. **(E)** Representative GFP⁻ and GFP⁺ gating regions visualised using FITC-A versus PE-A to facilitate discrimination from autofluorescent events. A gap was left between GFP⁻ and GFP⁺ gates. **(F)** GFP⁺ cell frequency across liver, heart, kidney, lung, spleen and muscle single-cell suspensions. Each dot represents an individual mouse/sample, with mean ± SEM indicated. GFP⁺ frequency was calculated as a percentage of singlet events using histogram-based gating.

Before analysing ASO-mediated *Malat1* knockdown, we performed RT-qPCR assays using standard-curve analysis and amplicon-size validation for *Malat1* and the reference genes *Ap3d1* and *Rpl13a* (Supplementary Figure S1A–C). *Malat1* was detectable in tissue-derived single-cell suspensions and whole-tissue RNA using both reference-gene normalisation approaches, supporting its use as a pharmacodynamic readout in tissue-derived samples (Supplementary Figure S1D–G). These baseline measurements were used to confirm assay detectability and normalisation performance, rather than to compare absolute *Malat1* abundance between tissues.

Freshly prepared tissue-derived single-cell suspensions were then analysed by flow cytometry. Cells were first gated using forward and side scatter to exclude debris, followed by singlet discrimination (Figure 2B–C). GFP⁺ cell frequency was quantified using histogram-based gating and expressed as a percentage of singlet events (Figure 2D). In parallel, conservative GFP⁺ and GFP⁻ gates were applied for FACS isolation, with a gap between gates to avoid collection of cells with intermediate fluorescence (Figure 2E). Additional representative gating examples across liver, lung, kidney, spleen and muscle confirmed that the same strategy could be applied across tissue-derived suspensions (Supplementary Figure S2).

GFP⁺ cells were detectable across all analysed tissues but represented a minority of singlet events, with higher frequencies in lung (8.72%), skeletal muscle (6.18%) and heart (5.1%) and lower frequencies in kidney (0.86%), liver (0.54%) and spleen (0.13%) (Figure 2F). Short-term culture of tissue-derived single-cell suspensions further confirmed the presence of GFP⁺ cells in multiple organ-derived preparations, although GFP⁺ cell abundance and morphology varied across tissues (Supplementary Figure S3). Together, these data established a workflow for quantifying GFP⁺ cell abundance and isolating GFP⁺ and GFP⁻ reporter-defined cell populations for downstream analysis of ASO-mediated *Malat1* knockdown.

### Whole-tissue *Malat1* knockdown is tissue- and conjugate-dependent

Systemic ASO activity was first evaluated by assessing *Malat1* expression across multiple tissues 5.5 days after a single subcutaneous administration of unconjugated or conjugated *Malat1*-targeting gapmer ASOs (Figure 3). ASO-mediated *Malat1* reduction varied substantially between tissues and conjugate architectures. The unconjugated ASO showed its strongest whole-tissue activity in the liver, with an approximately 30% reduction in *Malat1* expression (P < 0.05). In comparison, conjugated ASOs generally showed a broader distribution of knockdown across extrahepatic tissues, although both the magnitude and tissue profile differed between conjugates.

**Figure 3.**
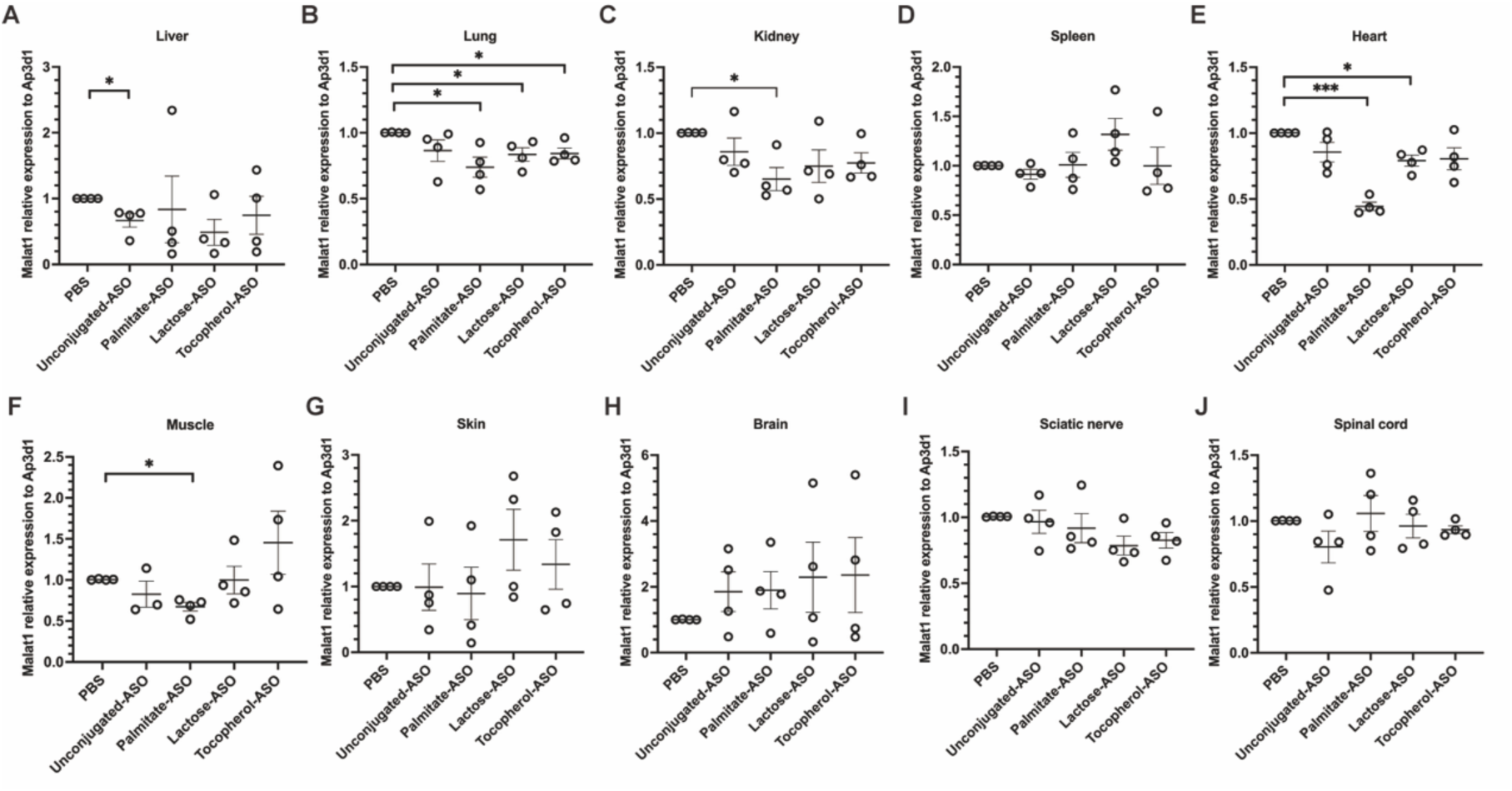
Whole-tissue *Malat1* expression following ASO treatment *in vivo*. **(A–J)** Relative Malat1 expression in liver (A), lung (B), kidney (C), spleen (D), heart (E), muscle (F), skin (G), brain (H), sciatic nerve (I) and spinal cord (J) 5.5 days after subcutaneous administration of PBS, unconjugated ASO, palmitate-ASO, lactose-ASO or tocopherol-ASO. Expression values were normalised to *Ap3d1* and further normalised to the PBS-treated group within the same tissue. Each point represents an individual mouse; bars indicate mean ± SEM. Statistical analysis was performed by one-sample t-test comparing each ASO-treated group with the PBS-normalised value of 1. *P < 0.05, **P < 0.01, ***P < 0.001.

Among the conjugates, palmitate-ASO produced the broadest whole-tissue knockdown, with significant *Malat1* reductions detected in lung, kidney, heart, and skeletal muscle. The strongest effect was observed in heart, where *Malat1* expression was reduced by approximately 60% (P < 0.001), while significant knockdown was also detected in kidney (∼35% reduction; P < 0.05), lung and skeletal muscle (∼30% reduction; P < 0.05 in both). Lactose-ASO displayed a more restricted profile, with significant reductions observed in lung and heart (∼20%; P < 0.05 in both), whereas tocopherol-ASO showed only a modest but significant reduction in lung (∼20%; P < 0.05). No significant *Malat1* reduction was detected in spleen, skin, brain, sciatic nerve, or spinal cord. Collectively, these findings demonstrate that conjugation did not uniformly increase ASO activity but instead generated distinct tissue-dependent pharmacodynamic profiles, with palmitate conjugation providing the broadest extrahepatic activity under the conditions tested.

### Cell-resolved analysis reveals tissue-dependent patterns of pharmacodynamic ASO activity

To determine whether whole-tissue *Malat1* reduction reflected efficient ASO knockdown within specific cell populations, isolated cells were separated into PDGFRα-GFP⁺ and GFP⁻ populations by FACS. *Malat1* expression was subsequently quantified within each cell population relative to the corresponding PBS control group (Figure 4). Cell-resolved pharmacodynamic activity varied substantially between organs and did not consistently reflect the patterns observed in whole-tissue samples. The most pronounced differences between whole-tissue and cell-resolved knockdown were observed in heart and skeletal muscle. In heart, the palmitate-ASO produced the strongest whole-tissue *Malat1* reduction (∼60%; P < 0.001), with a lower but significant reduction observed for lactose-ASO (∼20%; P < 0.05). In contrast, no significant *Malat1* reduction was detected for any ASOs in the GFP⁻ population, while only the unconjugated ASO presented a small while still significant reduction in GFP⁺ cells (∼20%; P < 0.05). Thus, the pronounced whole-heart knockdown activity of palmitate-ASO was not recapitulated in either GFP⁺ or GFP^-^cell population, highlighting a marked discrepancy between whole-tissue and cell-population-resolved pharmacodynamic measurements.

**Figure 4.**
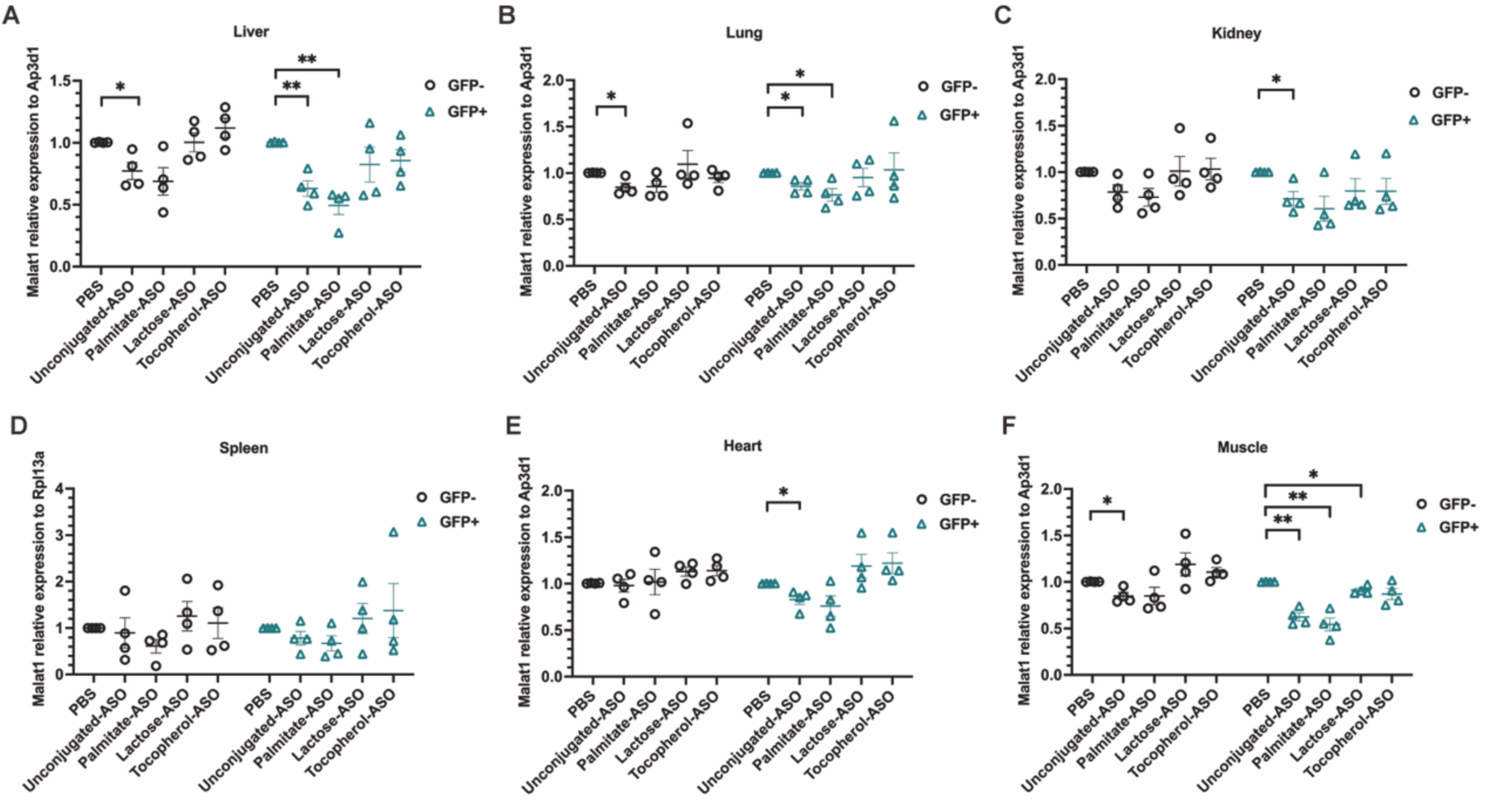
Cell-population-resolved *Malat1* target engagement in GFP⁺ and GFP⁻ population following ASO treatment. **(A–F)** Malat1 expression in GFP⁻ and GFP⁺ fractions sorted from liver (A), lung (B), kidney (C), spleen (D), heart (E) and muscle (F) after subcutaneous administration of PBS, unconjugated ASO, palmitate-ASO, lactose-ASO or tocopherol-ASO. Expression was normalised to *Ap3d1*, except for spleen samples, which were normalised to *Rpl13a*, and further normalised to the corresponding PBS-treated group within each sorted population. Each point represents an individual mouse; bars indicate mean ± SEM. Statistical analysis was performed by one-sample t-test against the PBS-normalised value of 1. *P < 0.05, **P < 0.01, ***P < 0.001

In skeletal muscle, the whole-tissue knockdown activity was comparatively modest, with palmitate-ASO reducing *Malat1* by approximately 30% (P < 0.05). Following cell sorting, however, the knockdown activity was substantially more pronounced within the GFP⁺ population, where unconjugated ASO reduced *Malat1* by approximately 40% (P < 0.01) and further reduction by palmitate-ASOs by 50% (P < 0.01). A very modest reduction was also observed for lactose-ASO (∼10%; P < 0.05). In comparison, *Malat1* reduction in the GFP⁻ population was limited, with only the unconjugated ASO demonstrating modest reduction (∼15%; P < 0.05). These findings indicate that efficient target gene knockdown in skeletal muscle can be achieved by palmitate-ASO within PDGFRα^+^ stromal populations, including FAPs. However, this knockdown activity could be underestimated by whole-tissue analysis alone.

In the lung, despite containing the highest proportion of GFP⁺ cells (8.72%) among the organs analysed, the unconjugated ASO produced comparable *Malat1* reduction in GFP⁺ and GFP⁻ populations (∼15% each; P < 0.05 in both), while palmitate-ASO showed modest knockdown activity in GFP⁺ cells (∼25% reduction; P < 0.05) comparable to the knockdown activity achieved in the whole-tissue assay. There was no effect in the GFP⁻ cell population. The relatively similar knockdown activity between GFP⁺ population and whole-tissue suggest that efficient ASO knockdown activity in lung was likely associated with the PDGFRα-reporter-positive cell population.

Additional tissue-specific patterns were observed in liver and kidney. In liver, unconjugated and palmitate-ASOs showed pronounced activity in GFP⁺ cells, reducing *Malat1* by approximately 40% and 50%, respectively (P < 0.01 for both). In GFP⁻ population, the knockdown activity was more moderate and reached statistical significance only for the unconjugated ASO (∼20% reduction; P < 0.05). Kidney showed comparable trends of knockdown activities of the unconjugated and palmitate-ASOs as presented in the liver, with more profound *Malat1* reduction in GFP^+^ cell populations (∼30% reduction for unconjugated ASO, P < 0.05; ∼40% reduction for palmitate-ASO, although not statistically significant). The reduction levels in GFP^+^ cells are more comparable to the whole-kidney knockdown activity observed for palmitate-ASO (∼35% reduction; P < 0.05).

Direct paired comparison of GFP⁺ and GFP⁻ populations within each treatment group further revealed tissue- and treatment-dependent differences in pharmacodynamic response (Supplementary Figure 4). Significantly greater *Malat1* knockdown was observed in GFP⁺ than GFP⁻ populations in heart following unconjugated ASO (P < 0.01) and palmitate-ASO (P < 0.05), in skeletal muscle following unconjugated ASO (P < 0.05), palmitate-ASO (P < 0.01) and tocopherol-ASO (P < 0.01), and in kidney following lactose-ASO (P < 0.05) and tocopherol-ASO (P < 0.01). No significant GFP⁺ versus GFP⁻ differences were detected for any treatment in liver, lung or spleen.

Collectively, these findings demonstrate that ASO pharmacodynamic activity differed across tissues and cell populations, and that whole-tissue responses did not consistently reflect activity within reporter-defined populations. The most notable discrepancies were observed in heart and skeletal muscle, with palmitate-ASO showing pronounced whole-tissue activity in heart and enhanced activity within the GFP⁺ skeletal-muscle population. Direct GFP⁺ versus GFP⁻ comparisons further identified tissue- and conjugate architecture-dependent population differences in heart, skeletal muscle and kidney.

### Preliminary safety assessment reveals tocopherol-ASO-associated toxicity

Body weight monitoring, macroscopic organ examination and plasma biochemical analysis were used to assess short-term toxicity following a single subcutaneous ASO administration in 8-week-old mice (2 male and 2 female mice/group). In most groups, body weight increased over the 5.5-day experimental period in both male and female mice in line with the body-weight gain during normal growth. However, male mice treated with tocopherol-ASO showed 4.56% reduction in body weight, whereas no comparable change was observed in tocopherol-ASO-treated females (Figure 5A–B).

**Figure 5.**
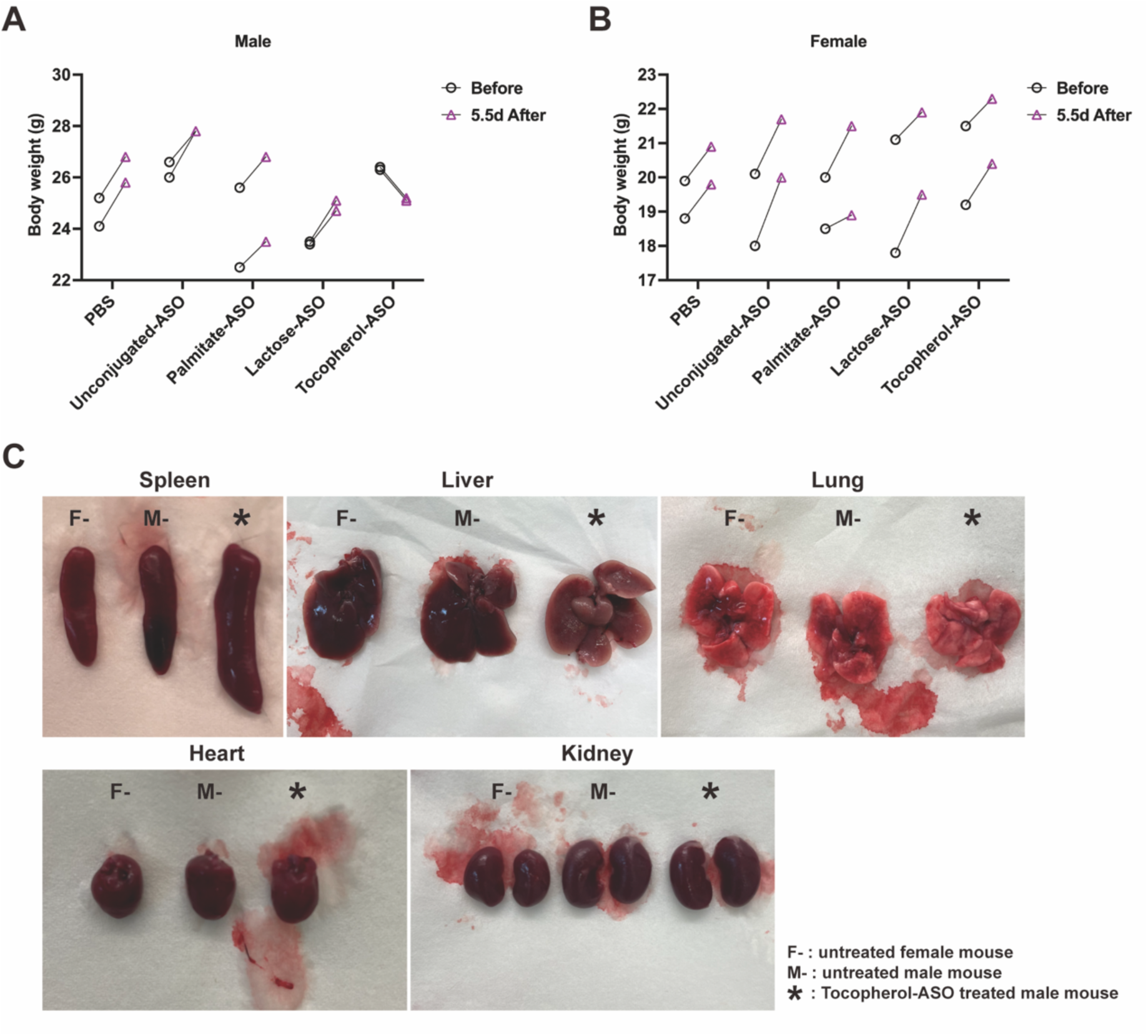
Short-term tolerability assessment following ASO administration. **(A–B)** Body weight measurements before treatment and 5.5 days after subcutaneous administration of PBS, unconjugated ASO, palmitate-ASO, lactose-ASO or tocopherol-ASO in male (A) and female (B) mice. Paired points represent individual animals before and after treatment. **(C)** Representative macroscopic images of spleen, liver, lung, heart and kidney collected at endpoint. Organs from a tocopherol-ASO-treated male mouse are shown alongside age-matched untreated female and male control mice.

Macroscopic examination revealed notable spleen enlargement in tocopherol-ASO- treated mice, in both male and female, compared with age-matched untreated controls (Figure 5C and Supplementary Figure S5). Other examined organs, including liver, lung, heart and kidney, did not show any obvious gross abnormalities in the same group of mice. No gross organ abnormalities were detected in mice from any other unconjugated, palmitate- or lactose-ASOs treated groups.

In the tocopherol-ASO group, ALT values were elevated in the two male mice (200 and 774 U/L), whereas normal values in females (52 and 32 U/L). Together with body-weight loss in male mice and spleen enlargement observed in both sexes, these findings suggest potential acute tocopherol-ASO-associated toxicity under the conditions tested, with the most pronounced effects observed in males.

**Table 1.** Plasma biochemical parameters following ASO administration. Plasma biochemical measurements obtained 5.5 days after subcutaneous administration of PBS, unconjugated ASO, palmitate-ASO, glucose-ASO/lactose-ASO or tocopherol-ASO. Mice were grouped by treatment and sex. Alanine aminotransferase (ALT), gamma-glutamyl transferase (GGT), total bilirubin (TBIL), albumin (ALB), creatinine (CRE), blood urea nitrogen (BUN) and uric acid (UA) †, values potentially affected by haemolysis interference.

| **Treatment** | **Sex** | **ALT** | **GGT** | **TBIL** | **ALB** | **CRE** | **BUN** | **UA** | **Hemolysis** |
|---|---|---|---|---|---|---|---|---|---|
|  |  | **U/L** | **U/L** | **mg/dL** | **g/L** | **mg/dL** | **mg/dL** | **mg/dL** |  |
| PBS | Male | 47 | 25 | 3.5 | 37 | 0.46 | 32.2 | 4.2 | Normal |
|  | Male | 59 | 87 | 5.7 | 43 | 0.36 | 35.7 | 4 | Mild |
|  | Female | 49 | 17 | 3.4 | 38 | 0.3 | 26.6 | 3.9 | Normal |
|  | Female | 33 | <10 | 2.8 | 39 | 0.31 | 35.3 | 4.5 | Mild |
| Unconjugated-ASO | Male | 52 | **340** <sup>†</sup> | **16.3** <sup>†</sup> | **>60** <sup>†</sup> | 0.61 | 34.9 | 3.6 | **Serious** |
|  | Male | 83 | 44 | 3.7 | 37 | 0.45 | 16.5 | 4.8 | Mild |
|  | Female | 42 | <10 | 1.5 | 31 | 0.35 | 22 | 5.4 | Normal |
|  | Female | 41 | <10 | 2.7 | 39 | 0.37 | 24.4 | 4.3 | Normal |
| Palmitate-ASO | Male | 28 | 98 | 5.7 | 42 | 0.61 | 29 | 5.0 | Mild |
|  | Male | 35 | 18 | 4.2 | 35 | 0.51 | 38.4 | 5.6 | Mild |
|  | Female | 65 | **333**<sup>†</sup> | **15.5** <sup>†</sup> | **>60** <sup>†</sup> | 0.41 | 24.3 | 3.7 | **Serious** |
|  | Female | 34 | <10 | 1.6 | 36 | 0.28 | 20.7 | 4.5 | Normal |
| Lactose-ASO | Male | 72 | 93 | 6.4 | 48 | 0.64 | 23.9 | 3.1 | Mild |
|  | Male | 32 | 14 | 3 | 38 | 0.5 | 29.2 | 5.1 | Mild |
|  | Female | 65 | **321**<sup>†</sup> | **17.8** <sup>†</sup> | **>60** <sup>†</sup> | 0.37 | 28.5 | 3.4 | **Serious** |
|  | Female | 28 | 44 | 5.2 | 43 | 0.33 | 20.9 | 5.6 | Mild |
| Tocopherol-ASO | Male | **200** | 22 | 3.3 | 25 | 0.57 | 17.3 | 6.2 | Mild |
|  | Male | **774** | 10 | 2.5 | 20 | 0.72 | 19 | 5.6 | Normal |
|  | Female | 52 | 20 | 4.4 | 37 | 0.59 | 29.5 | 7.3 | Mild |
|  | Female | 32 | <10 | 1.6 | 38 | 0.37 | 22.6 | 4.9 | Normal |

† Hemolysis interference

## DISCUSSION

This study used *Malat1* knockdown as a functional pharmacodynamic readout to compare the whole-tissue and cell-population-resolved activity of unconjugated and chemically conjugated ASOs *in vivo*.^23, 24^ By combining whole-tissue RT-qPCR with FACS isolation of PDGFRα-eGFP reporter-defined populations, we identified two principal findings. First, ASO activity depended strongly on tissue context and conjugate architecture. Second, whole-tissue knockdown did not consistently predict pharmacodynamic activity within the corresponding GFP^+^ and GFP^-^reporter-defined populations. These results show that organ-level pharmacodynamic measurement and cell-resolved analyses provide complementary, rather than interchangeable information about productive ASO activity.

At the whole-tissue level, conjugation did not produce a uniform increase in target reduction. Instead, each conjugate generated a distinct activity profile. The Palmitate-ASO showed the broadest pattern of significant extrahepatic activity, with the strongest effect observed in heart. In liver, lactose-ASO produced the largest numerical reduction, whereas the unconjugated ASO showed the clearest statistically significant effect. Lactose- and tocopherol-ASOs otherwise displayed more restricted tissue activity. These findings indicate that ASO chemical conjugation can reshape functional activity *in vivo*, generating tissue-specific pharmacodynamic profiles rather than a uniform increase in target reduction across organs. This is consistent with the broader understanding that conjugation can alter ASO pharmacokinetics, plasma-protein interactions and tissue distribution without necessarily producing a proportional increase in productive intracellular activity.^5–7, 19, 34, 35^

The activity profile of palmitate-ASO is consistent with previous reports that fatty-acid conjugation can enhance the potency of phosphorothioate ASOs in muscle tissues.^19, 34, 35^ Increasing fatty-acid chain length strengthens plasma-protein binding, and palmitate conjugation has been shown to increase plasma exposure and ASO potency in mouse muscle.^35^ Mechanistic studies further showed that palmitate increases ASO exposure in heart and skeletal muscle, although much of the ASO accumulates in the interstitial space rather than intracellularly.^19^ These observations distinguish tissue exposure from cellular uptake and productive intracellular activity in specific cell populations of interest. That distinction is directly relevant here: palmitate-ASO produced approximately 60% *Malat1* reduction in whole heart, yet no significant reduction relative to PBS was detected in either the PDGFRα⁺/GFP⁺ or or PDGFRα⁻/GFP⁻ sorted population. The whole-heart response therefore cannot be inferred from reporter-defined cell activity alone.

Comparison of palmitate- and tocopherol-ASOs further indicates that lipophilicity alone does not determine productive activity. Although both conjugates contain lipophilic moieties, palmitate-ASO showed substantially broader activity, including significant whole-tissue knockdown in heart and skeletal muscle, whereas tocopherol- ASO did not. Previous work has shown that palmitate and tocopherol conjugates differ in their plasma-protein associations and pharmacological behaviour, with palmitate associating predominantly with albumin and tocopherol showing greater association with lipoproteins.^34^ Tocopherol-ASO pharmacodynamic activity has also been reported to depend on administration route, with greater target knockdown following intravenous than subcutaneous dosing.^34^ The subcutaneous route used here may therefore have contributed to its limited activity.

These comparisons must nevertheless be interpreted at the level of the complete conjugate architecture. The palmitate-ASO used here contains a C6 linker matching previously reported constructs,^19, 34, 35^ whereas the tocopherol-ASO contains an ethylene glycol linker and lacks an additional phosphate-containing linker feature present in some earlier designs. Lactose-ASO is chemically distinct and was generated by BCN-SPAAC, which introduced a comparatively bulky BCN-derived cycloadduct between the carbohydrate and the ASO. Consequently, differences among the compounds may reflect ligand identity, linker composition, charge, stability, protein binding, cellular uptake or intracellular processing. BCN-containing linker architecture and linker composition can influence conjugation performance and functional activity in other ASO conjugate formats.^36^ The restricted activity of lactose-ASO should therefore not be interpreted as an intrinsic property of lactose, just as the different behaviours of the two lipophilic conjugates cannot be attributed solely to palmitate or tocopherol. Matched-linker controls and systematic linker series will be needed to separate these effects.

Cell-resolved analysis revealed additional pharmacodynamic heterogeneity that was not apparent from whole-tissue measurements. This agrees with the work of Shin *et al.*, who showed that ASO silencing in lung differs among cell populations and that bulk measurements can obscure strong activity in individual compartments.^24^ Following intratracheal delivery, they observed robust silencing in PDGFRα/CD140a-positive lung fibroblasts and endothelial cells, while tissue ASO concentration did not directly predict silencing efficacy.^24^ Although that study used local administration and a different ASO chemistry, it established the importance of distinguishing tissue accumulation from productive activity and of resolving pharmacodynamic effects at the cellular level. The present study extends this approach to systemically administered ASO conjugates across multiple tissues.

Skeletal muscle provided the clearest example of whole-tissue analysis underestimating activity in a reporter-defined population. Palmitate-ASO reduced whole-muscle *Malat1* by approximately 30%, but the reduction in PDGFRα⁺ cells was approximately 50% and was greater than that observed in PDGFRα⁻ cells. The unconjugated ASO also showed stronger activity in PDGFRα⁺ than PDGFRα⁻ cells, and lactose-ASO produced a smaller but significant effect in PDGFRα⁺ cells. Because PDGFRα⁺ cells represented only 6.18% of singlet events, pronounced activity in this minority population would be expected to contribute only modestly to total tissue RNA. Previous studies showed that palmitate conjugation increases ASO exposure and potency in skeletal muscle.^19, 35^ Early histological analysis of quadriceps further showed ASO staining concentrated along intercellular boundaries, consistent with localisation in the interstitial space or plasma membrane, with punctate intracellular staining becoming apparent at later time points.^19^ Our findings extend these previous tissue-level observations by providing cell-resolved evidence of preferential productive pharmacodynamic activity within the PDGFRα-reporter-positive skeletal-muscle stromal compartment, which includes FAPs, particularly following palmitate-ASO treatment. This cell-resolved pattern may therefore be relevant to fibroblast-directed therapeutic strategies in muscular dystrophies, including collagen VI-related disease where both splice switching ASO, or silencing approaches have experimentally demonstrated to be efficacious in cellular models.^32, 37–40^

Liver and kidney showed related but less definitive patterns. In liver, unconjugated and palmitate-ASOs reduced *Malat1* by approximately 40% and 50%, respectively, in PDGFRα⁺ cells, whereas activity in PDGFRα⁻ cells was more modest and reached significance only for the unconjugated ASO. However, direct PDGFRα⁺ versus PDGFRα⁻ comparisons were not significant in liver. In kidney, unconjugated and palmitate-ASOs also showed numerically greater reductions in PDGFRα⁺ cells, but the palmitate effect did not reach significance in that population, and paired comparisons showed significantly greater knockdown in PDGFRα⁺ than PDGFRα⁻ cells was instead detected for lactose- and tocopherol-ASOs. These findings suggest cell-population-dependent pharmacodynamic activity, but the small PDGFRα⁺ frequencies and limited sample size require cautious interpretation.

Heart illustrated the converse relationship. Palmitate-ASO produced the strongest whole-tissue response in the study, yet neither PDGFRα⁺ nor PDGFRα⁻ populations showed significant reduction relative to their respective PBS controls. A paired analysis nevertheless detected greater knockdown in PDGFRα⁺ than PDGFRα⁻ cells after palmitate-ASO treatment. These results are not contradictory: the paired comparison assesses the relative response of the PDGFRα⁺ and PDGFRα⁻ sorted populations, whereas comparisons with PBS assess whether knockdown within each population is individually distinguishable from control. As PDGFRα⁺ cells represented only 5.1% of singlet events, they are unlikely to account for the approximately 60% whole-heart reduction. Previous studies showed that palmitate conjugation increases cardiac ASO exposure and potency. Early histological analysis following a single subcutaneous dose showed diffuse or intercellular ASO staining, consistent with localisation in the interstitial space, with punctate intracellular staining becoming apparent by 24 h.^19, 35^ These previous observations suggest that palmitate-ASO initially distributes largely within the cardiac interstitial compartment, but they do not identify the specific cell populations that subsequently take up the ASO and mediate productive target engagement. The strong whole-heart response observed here could therefore reflect productive uptake across multiple cardiac cell populations, only a subset of which may be represented by the PDGFRα-reporter-positive compartment. The adult heart contains cardiomyocytes, endothelial and mural cells, immune cells and heterogeneous stromal populations, including multiple fibroblast subsets.^41^ Accordingly, the present data cannot determine which of these populations contributes most to the whole-heart response. Future studies using expanded marker panels, single-cell or single-nucleus approaches, and direct ASO quantification within defined cardiac cell populations will be needed to identify the cellular contributors to the whole-heart response.

Lung showed a third pattern. It contained the highest PDGFRα⁺ frequency among the tissues examined (8.72%). The unconjugated ASO reduced *Malat1* similarly in PDGFRα⁺ and PDGFRα⁻ cells, whereas palmitate-ASO produced significant knockdown in PDGFRα⁺ cells but not in PDGFRα⁻ cells. However, direct paired comparisons between the two populations were not significant for any treatment. The lung data therefore suggest activity within the PDGFRα-reporter-positive compartment without establishing preferential activity relative to PDGFRα⁻ cells. The difference from the marked fibroblast susceptibility reported after intratracheal LNA-gapmer administration,^24^ may reflect route of administration, systemic exposure, ASO chemistry or the cellular populations recovered by each method. More broadly, the heart, skeletal-muscle and lung results demonstrate that whole-tissue activity, population-specific activity and preferential activity between populations are related but distinct pharmacodynamic measures.

The PDGFRα-eGFP model demonstrates the utility of reporter-based approaches for systematically evaluating ASO conjugate activity across multiple organs, providing a valuable translational framework for investigating productive ASO activity within defined cellular populations. By enabling the isolation of reporter-positive and reporter-negative populations using a common fluorescence-based sorting strategy, this model provides a practical complement to approaches requiring extensive cell-type-specific marker panels or single-cell analysis. Unlike studies restricted to individual organs or administration routes, this framework allows pharmacodynamic responses to different ASO conjugate architectures to be compared across tissues following systemic administration. Thus, it establishes an experimental approach for determining whether whole-tissue ASO activity reflects productive activity within a defined cellular compartment.

Moreover, the biological relevance of the PDGFRα-associated compartment further strengthens this approach. PDGFRα^+^ stromal populations contribute to tissue remodelling and fibrosis across multiple organs, while skeletal-muscle PDGFRα^+^ fibro-adipogenic progenitors regulate regeneration and can contribute to fibrosis and disease progression in muscular dystrophies.^42–44^ The present study used *Malat1* solely as a model target, but the workflow could be applied to therapeutic ASOs directed against transcripts involved in extracellular-matrix production, fibrotic signalling, inflammatory crosstalk or fibroblast activation. Cell-resolved pharmacodynamic assessment may be particularly informative when the disease-relevant population represents only a small fraction of total tissue, as in skeletal muscle, where whole-tissue measurements may not reflect activity within the target population. Collagen VI-related muscular dystrophy provides a specific example, in which allele specific silencing of dominant gain of function mutation, or splice correction of both dominant and recessive mutant *COL6* transcripts in fibroblast-lineage cells represent a potential therapeutic target.^32^ More broadly, the same approach could be applied to other genetic disorders and acquired fibrotic diseases involving stromal-cell dysfunction, extracellular-matrix dysregulation or pathological remodelling.

The study also has several limitations related to the experimental model and study design. PDGFRα-eGFP identifies tissue-dependent reporter-positive compartments rather than a single uniform fibroblast population, and PDGFRα⁻ fractions remain heterogeneous. The use of a single dose, one 5.5-day endpoint and modest group sizes limits assessment of dose-response relationships, temporal pharmacology and sex-specific effects. ASO concentrations were not measured in plasma, whole tissues or sorted cells, so the relative contributions of systemic exposure, tissue accumulation, cellular uptake and intracellular trafficking cannot be resolved. Finally, because ligand and linker differed simultaneously, matched-linker controls will ideally be required to assign pharmacological effects to individual architectural features, which nevertheless was not the specific focus of our study.

The preliminary safety findings further distinguished the conjugates. Tocopherol-ASO was associated with body-weight loss and ALT elevations in the two male mice, together with spleen enlargement in both sexes, whereas comparable signals were not observed for the other ASO groups under the same conditions. These preliminary safety observations will need to be addressed with future more specific studies. The small number of animals, short follow-up and haemolysis affecting several plasma measurements preclude us from making firm conclusions about incidence, mechanism or sex dependence of the observed safety signals. The results nevertheless support more detailed dose-ranging studies with larger cohorts, longitudinal clinical chemistry, histopathology and exposure measurements before further evaluation of this tocopherol architecture.

In summary, ASO conjugation reshapes pharmacodynamic activity in a tissue- and cell-population-dependent manner that cannot be fully captured by conventional whole-tissue measurements. Palmitate conjugation not only produced the broadest extrahepatic activity but also enhanced productive target engagement within PDGFRα^+^ skeletal-muscle stromal cells, a population that includes fibro-adipogenic progenitors and represents a potential therapeutic target in muscular dystrophies and fibrotic disorders. These findings establish the PDGFRα-eGFP model as a practical system for evaluating cell-specific ASO activity in vivo and demonstrate the value of combining chemical delivery strategies with cell-resolved pharmacodynamic analysis. Such approaches should facilitate the development of next-generation ASO conjugates capable of reaching disease-relevant cellular targets rather than simply increasing tissue exposure, thereby improving the translational potential of cell-specific oligonucleotide therapeutics.

## MATERIALS AND METHODS

### Oligonucleotide synthesis, conjugation, and deprotection

Solid-phase synthesis of antisense oligonucleotides (ASOs) was performed using a K&A H-16 synthesizer (Sierra BioSystems). DNA and 2′-O-MOE phosphoramidites were obtained from ChemGenes, and all cytidine residues in the DNA and 2′-O-MOE regions were incorporated as 5-methylcytidine (5-Me-C). Lipid-conjugate phosphoramidites, including 5′-Palmitate-C6 amidite and Tocopherol TEG phosphoramidite, were obtained from ChemGenes, while the cyclic alkyne phosphoramidite bicyclo[6.1.0]nonyne (BCN) was purchased as 5′-Click-Easy™ BCN CE-Phosphoramidite I (LGC Biosearch Technologies) to enable Cu-free click conjugation. All phosphoramidites were dissolved in anhydrous acetonitrile (ACN) to a final concentration of 0.1 M, except for the Tocopherol TEG phosphoramidite, which was prepared at 0.05 M, and the Palmitate-C6 phosphoramidite, which was prepared in a 3:1 (v/v) mixture of dichloromethane (DCM) and ACN.

Detritylation was performed using 3% (v/v) trichloroacetic acid in DCM (Sigma-Aldrich). A 0.3 M solution of Hyacinth 5-benzylmercaptotetrazole (BMT; EMP, supplied by Atom Scientific Ltd) in anhydrous acetonitrile was used as the activator during phosphoramidite coupling. Sulfurization was carried out using 0.1 M DDTT (ChemGenes) in anhydrous pyridine (Sigma-Aldrich), while oxidation was performed using Hyacinth Oxidizer (EMP, supplied by Atom Scientific Ltd), consisting of 0.05 M iodine in a 9:1 (v/v) mixture of pyridine and water. Remaining 5′-hydroxyl groups were capped using Capping A (EMP, supplied by Atom Scientific Ltd), comprising an 8:1:1 (v/v/v) mixture of tetrahydrofuran (THF), lutidine, and acetic anhydride, in combination with Capping B (EMP, supplied by Atom Scientific Ltd), a 16% (w/v) solution of *N*-methylimidazole in THF.

Following synthesis, Tocopherol-bearing oligonucleotides still bound to the solid phase were treated with 20% (v/v) Diethylamine in anhydrous ACN for 5 min.

Deprotection and cleavage from the solid support were achieved using 30% aqueous ammonium hydroxide (NH_4_OH; Sigma-Aldrich) at 55°C for 18 h. Afterwards, the solution was allowed to cool to room temperature before the remaining ammonia was removed under reduced pressure using a Speedvac concentrator. Following the deprotection procedure, the oligonucleotides were precipitated in Ethanol puriss (Sigma-Aldrich) and in a 0.3 M Sodium acetate buffer (pH 5.2; ThermoFisher) at - 20 °C for 18 h. The precipitated oligonucleotide formed a pellet after the samples were centrifuged at 4550 rpm for 30 min at 4 °C. The resulting pellet was washed twice with 20 mL ice cold 70% ethanol, followed by centrifugation at 4550 rpm for 30 min at 4 °C. Finally, the pellets were dried under reduced pressure, and oligonucleotides were resuspended in RNase-free water (ThermoFisher). If required, additional desalting was achieved using 0.5 mL 3 kDa Amicon Ultra centrifugal filter units (Merck).

Liquid chromatography–mass spectrometry (LC-MS) analysis was performed using an ACQUITY Premier UPLC system coupled to a single quadrupole mass detector (SQD2; Waters). Purified oligonucleotides were diluted to a final concentration of 20 µM in water, and 50 µL aliquots were analysed in negative ionisation mode. Separation was achieved using an ACQUITY UPLC BEH C18 column with VanGuard (130 Å, 1.7 µm, 2.1 × 50 mm; Waters). Eluent A consisted of 8.6 mM triethylamine (TEA) and 100 mM 1,1,1,3,3,3-hexafluoroisopropanol (HFIP) in water, while eluent B was methanol. The flow rate was 0.5 mL min⁻¹, and the column temperature was maintained at 60 °C. The mass spectrometer was operated with a capillary voltage of 0.8 kV, a cone voltage of 20 V, and a desolvation temperature of 600 °C. Raw continuum data were deconvoluted to zero-charge mass spectra using MaxEnt1 software (Waters).

### Animal experiments and dosing

All animal procedures were conducted in accordance with the Animals (Scientific Procedures) Act 1986 and were approved by the UCL Animal Welfare and Ethical Review Body (AWERB) under project licence PP2611161. Six-to eight-week-old heterozygous PDGFRα-eGFP mice (B6.129S4-Pdgfra*^tm^*^11^*^(EGFP)Sor^*/J; The Jackson Laboratory, USA) were used, and genotyping was performed according to the protocol provided by The Jackson Laboratory.

Unconjugated ASO or ASO conjugates were administered by a single subcutaneous injection at 10.8 μmol/kg in PBS, with injection volumes adjusted according to body weight and ASO stock concentration. Injection volumes were approximately 80–200 μl. Control mice received 200 μl PBS by the same route. Body weight was recorded before dosing and at the experimental endpoint.

At 5.5 days post-dose, mice were placed under terminal anaesthesia, and blood was collected by cardiac puncture into K₂EDTA-coated tubes (Microvette, Germany) for plasma biochemical analysis. Following confirmation of death, tissues were collected for whole-tissue RNA extraction or immediate single-cell suspension preparation.

For whole-tissue RNA analysis, samples were placed immediately on dry ice and stored at −80°C until RNA extraction. For flow cytometry and fluorescence-activated cell sorting (FACS), selected samples were processed immediately after collection to generate single-cell suspensions.

### Whole-tissue RNA extraction

Whole-tissue RNA was extracted from liver, kidney, spleen, heart, lung, pooled hindlimb skeletal muscles, sciatic nerve, spinal cord, skin and brain. Tissue samples of approximately 10–50 mg were homogenised in 350-600μl RLT buffer supplemented with β-mercaptoethanol (10 μl per 1 ml RLT buffer) using ceramic bead tubes (1.4 mm, Fisherbrand X100 Tubes, Thermo Fisher Scientific) and a TissueLyser II (Qiagen). Homogenisation conditions were adjusted according to tissue type. Liver, spleen, brain and spinal cord samples were homogenised at 28 Hz for 1 min, incubated on ice for 1 min, and homogenised for a second cycle under the same conditions. Kidney, heart, lung, pooled hindlimb skeletal muscles, sciatic nerve and skin samples were homogenised at 30 Hz for 1 min, incubated on ice for 1 min, and homogenised for a total of three cycles. Lysates were clarified by centrifugation at 12000 rpm for 5 min at 4°C and passed through QIAshredder columns (Qiagen) prior to RNA isolation. RNA was extracted using the RNeasy Plus Mini Kit (Qiagen, Hilden, Germany, #74136) according to the manufacturer’s instructions.

### Single-cell suspension preparation

Single-cell suspensions were prepared from pooled hindlimb skeletal muscles, liver, kidney, spleen, heart and lung using tissue-specific enzyme-based dissociation protocols.

For pooled hindlimb skeletal muscle, tissues were washed in pre-cold PBS and mechanically minced into small fragments along the fibre axis. Tissues were subjected to a two-step enzymatic digestion. The first digestion was performed using collagenase II (250 U/mL, Sigma-Aldrich, USA) in PBS supplemented with CaCl₂ (final concentration 2.5 mM) at 37°C for 30 min with agitation (90 rpm). Following centrifugation at 156 × g for 5 min at 4°C, tissues were further digested in a second enzyme mixture containing collagenase D (1.5 U/mL, Roche, Switzerland) and dispase II (2.4 U/mL, Sigma-Aldrich, USA) in PBS supplemented with CaCl₂ (2.5 mM) under the same conditions for 1 h. Digestion was terminated using ice-cold PBS supplemented with 10% FBS and 4 mM ethylenediaminetetraacetic acid (EDTA, pH 7.9), after which cells were centrifuged at 300 × g for 5 min at 4°C.

For non-muscle tissues, including liver, kidney, spleen, heart and lung, samples were washed in pre-cold PBS, minced, and digested in RPMI-based buffer containing 1% fetal bovine serum (FBS), collagenase P (0.2 mg/mL, Sigma-Aldrich, USA), dispase II (0.8 mg/mL), DNase I (0.1 mg/mL, Sigma-Aldrich, USA), and CaCl₂ (0.25 mM) at 37°C for 20–30 min with agitation at 90 rpm, with digestion time adjusted according to tissue type. Supernatants were collected, and remaining tissue fragments were subjected to a second round of enzymatic digestion with additional mechanical dissociation under the same conditions for 20–30 min. Pooled cell suspensions were neutralised with stop solution consisting of RPMI supplemented with 10% FBS and 4 mM EDTA, followed by centrifugation at 233 × g for 5 min at 4°C. Cell pellets were treated with eBioscience 1× red blood cell Lysis Buffer (Thermo Fisher Scientific, USA; #00-4333-57), and lysis was stopped using the same stop solution. Cells were then centrifuged at 300 × g for 5 min at 4°C.

Following dissociation, all samples were resuspended in FACS buffer consisting of PBS supplemented with 2% FBS and 4 mM EDTA. Cell suspensions were sequentially filtered through 100 μm and 40 μm cell strainers (Corning), washed twice and resuspended in FACS buffer containing DNase I (1 μg/ml). Cell suspensions were maintained on ice until FACS.

### Short-term culture of tissue-derived cells

To assess the detectability of GFP⁺ cells following tissue dissociation, aliquots of freshly prepared single-cell suspensions were plated in Dulbecco’s Modified Eagle’s Medium (DMEM; Sigma-Aldrich, #D6429) supplemented with 10% FBS and cultured for 3 days under standard cell culture conditions. Adherent cells were imaged using an EVOS M5000 Imaging System (Invitrogen) to visualise brightfield morphology and GFP fluorescence.

### Flow cytometry and fluorescence-activated cell sorting

Tissue-derived single-cell suspensions were prepared as described above and maintained on ice before FACS. Where required, cell suspensions were filtered immediately before sorting to remove residual aggregates. Flow cytometry and FACS were performed on FACSAria III and FACSymphony S6 instruments (BD Biosciences, USA) according to standard operating procedures of the UCL GOS ICH Flow Cytometry Core Facility.

Cells were first gated using forward scatter area (FSC-A) and side scatter area (SSC-A) to exclude debris and select the main cell population. Singlet discrimination was performed using FSC-A versus FSC-W during cell sorting and FSC-A versus FSC-H for post-acquisition analysis. GFP fluorescence was detected in the FITC-A channel and plotted against PE-A to facilitate discrimination of GFP-expressing cells from autofluorescent events, without PE fluorophore staining. GFP⁺ and GFP⁻ populations were defined according to GFP fluorescence intensity. GFP gates were set using matched control samples processed on the same day to account for tissue-specific background fluorescence and day-to-day instrument variation. The same threshold was then applied to corresponding experimental samples within the same sorting batch. For GFP⁺ cell frequency quantification, histogram-based gating was used, with GFP⁺ events expressed as a percentage of singlet events. For cell sorting, conservative GFP⁺ and GFP⁻ gates were separated by a gap to exclude cells with intermediate GFP fluorescence.

Sorted GFP⁺ and GFP⁻ cell populations were collected into 5 ml round-bottom FACS tubes (Falcon) preloaded with FBS to support cell recovery and minimise cell loss. Collection volumes varied depending on the number of sorted cells and sorting duration, but matched tissue groups were collected under comparable conditions. Sorted cells were kept on ice and processed for downstream RNA extraction. Flow cytometry data were analysed using FlowJo software version 10 (BD Biosciences).

### RNA extraction from sorted cells

Sorted GFP⁺ and GFP⁻ cell suspensions were centrifuged at 3,000 rpm for 10 min at 4°C. Supernatants were carefully removed without disturbing the cell pellet. Total RNA was extracted from sorted cell pellets using the RNeasy Mini Kit (Qiagen, Hilden, Germany, #74106) according to the manufacturer’s instructions.

### Reverse transcription and quantitative PCR

RNA concentration and purity were assessed using a DS-11 Series spectrophotometer (DeNovix). One-step RT–qPCR was performed using the iTaq Universal Probes One-Step Kit (Bio-Rad, CA, USA, #1725141) on a StepOne Real-Time PCR System (Applied Biosystems). Reverse transcription was performed at 50°C for 10 min, followed by polymerase activation and initial denaturation at 95°C for 3 min. PCR amplification was then performed for 40 cycles of denaturation at 95°C for 10 s and annealing/extension at 60°C for 30 s.

Primers and probes were synthesised by Kaneka Eurogentec S.A. (Seraing, Belgium) as follow: *Malat1*(forward: 5′-TAC GGA TGT GGT GGT GAA GC-3′, reverse: 5′-TCT AGG AGC CGC TAT CTA TCC-3′, probe: 5′-[6FAM] TGG TGG AGC GAG AGC GAG TTG GT [BHQ1]-3′); *Ap3d1*(forward: 5′-CAG ATC GAC CTC TCA GTG CC-3′, reverse: 5′-TGT GAA GCT GGA GGA GCA GCG-3′, probe: 5′-[JOE] TTC TTG TCC TTC TCC AGC CG [BHQ1]-3′); and *Rpl13a* (forward: 5′-GCT GCT CTC AAG GTT GTT CG-3′, reverse: 5′-TCC GAT AGT GCA TCT TGG CC-3′, probe: 5′-[JOE] ACC AGG CAG TGA CAG CCA CTC TGG [BHQ1]-3′).

For assay validation, total RNA extracted from 3T3 mouse fibroblast cells was serially diluted and used to generate standard curves for *Malat1*, *Ap3d1* and *Rpl13a* RT-qPCR assays. 3T3 cells were maintained in DMEM supplemented with 10% FBS under standard cell culture conditions before RNA extraction.

### Plasma biochemistry

Blood was transferred to 1.5 ml microcentrifuge tubes and centrifuged at 8,000 rpm for 5 min to separate plasma. Plasma was collected and analysed using a dry chemistry analyser (FUJIFILM DRI-CHEM NX500, FUJIFILM Corporation, Japan) according to the manufacturer’s instructions. Plasma alanine aminotransferase (ALT), gamma-glutamyl transferase (GGT), total bilirubin (TBIL), albumin (ALB), creatinine (CRE), blood urea nitrogen (BUN) and uric acid (UA) were measured using the corresponding FUJI DRI-CHEM SLIDE reagents.

### Data processing and normalisation

For each mouse sample, RT-qPCR reactions were performed in technical triplicate. Technical replicate cycle threshold (Ct) values were reviewed for amplification consistency before analysis. Where all three technical replicates were concordant, the mean Ct of three wells was used. An SD of ≤0.2 Ct between technical replicates was used as the preferred quality-control criterion. Where one well deviated from the other two, the concordant duplicate was used for downstream analysis, provided that the difference between the two accepted wells did not exceed 0.5 Ct.

Baseline and fluorescence threshold settings were reviewed for each qPCR plate to ensure that Ct values were called within the exponential amplification phase. For each assay, threshold settings were kept consistent across comparable plates where amplification curves permitted. Where plate-specific adjustment was required, thresholds were set within the exponential phase, and treatment effects were interpreted after normalisation to plate-matched PBS controls.

*Malat1* expression was normalised to the corresponding reference gene using the delta Ct method. Relative expression was calculated using the delta-delta Ct method. To account for qPCR plate and experimental batch effects, treatment groups were normalised to PBS controls processed on the same plate or within the corresponding tissue-specific batch. Analyses were performed within each tissue and, for sorted samples, within each GFP-defined cell population. Relative expression values were therefore interpreted as tissue- and cell-population-specific *Malat1* target engagement rather than as comparisons of absolute *Malat1* abundance between tissues or between GFP-defined populations.

### Statistical analysis

Statistical analyses and data visualisation were performed using GraphPad Prism version 10.0. Each biological replicate represents an individual mouse, with four mice per treatment group unless otherwise stated. Data are presented as mean ± standard error of the mean (SEM). For relative expression data normalised to PBS controls, treatment groups were compared against a hypothetical value of 1.0 using a one-sample t-test. PBS controls were used as the normalisation reference and were not subjected to statistical testing against 1.0. *P* < 0.05 was considered statistically significant.

### Figure preparation

The experimental workflow schematic was created using BioRender.com. Chemical drawings were created using ChemDraw 25.5.0.

## DATA AVAILABILITY

Raw and processed RT-qPCR data, flow cytometry gating percentages and statistical test results supporting this study are provided in the Supplementary Data.

## ACKNOWLEDGEMENTS

We thank Fady Guirguis for sharing the protocols for preparing single-cell suspensions from skeletal muscle and liver. These protocols provided a useful starting point for the tissue dissociation workflows, which were subsequently adapted for the broader panel of tissues analysed in this study

## AUTHOR CONTRIBUTIONS

H.Z., J.F., M.W. and F.M. conceived the collaborative project. S.C. designed and performed the biological experiments, analysed and interpreted the data, and drafted the manuscript. J.F. contributed to ASO conjugate design and synthesis and drafted the manuscript. A.E. and P.C. performed fluorescence-activated cell sorting. H.Z., M.W. and F.M. reviewed and revised the manuscript. All authors reviewed and approved the final manuscript.

## SUPPLEMENTARY DATA

Supplementary Data are available online.

