## Supplementary Figures for "Cell-resolved pharmacodynamic profiling reveals tissue- and cell-population-specific activity of antisense oligonucleotide chemical conjugates *in vivo*"

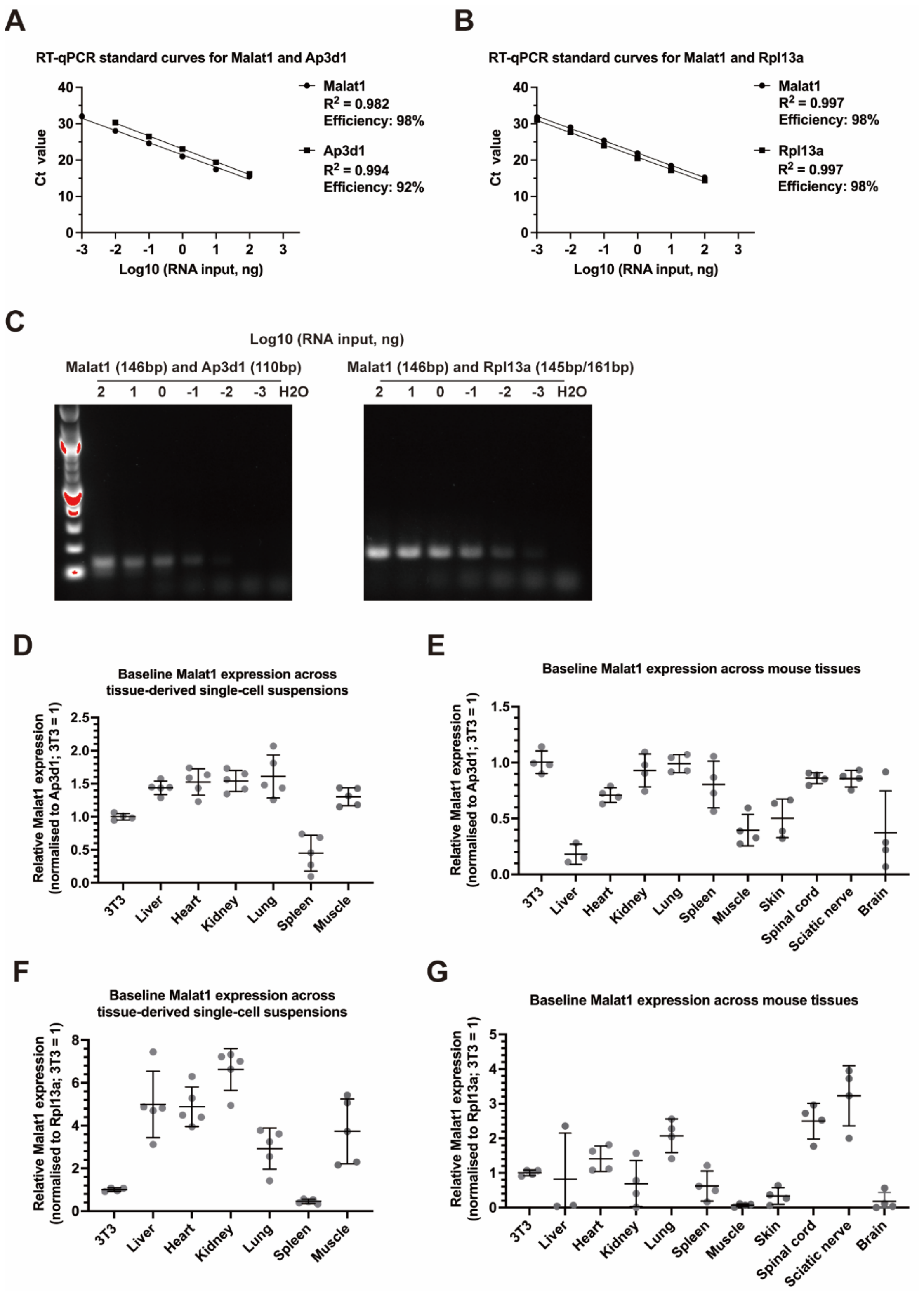


Supplementary Figure S1. RT-qPCR assay validation and baseline *Malat1* measurements in tissue-derived samples.

(A–B) RT-qPCR standard curves for Malat1 and reference genes generated using serial dilutions of 3T3 RNA. Standard curves are shown for *Malat1* with *Ap3d1* (A) and Malat1 with *Rpl13a* (B). $R^2$ values and amplification efficiencies are indicated.

(C) Agarose gel electrophoresis of RT-qPCR amplicons across the dilution series.

Expected product sizes are indicated for Malat1, *Ap3d1* and *Rpl13a*. $H_2O$ indicates the no-template control.

(D–E) Malat1 expression in untreated tissue-derived single-cell suspensions (D) and whole-tissue RNA samples (E), normalised to *Ap3d1* and expressed relative to 3T3 RNA.

(F–G) Malat1 expression in untreated tissue-derived single-cell suspensions (F) and whole-tissue RNA samples (G), normalised to *Rpl13a* and expressed relative to 3T3 RNA.

Each dot represents an individual mouse/sample, with mean ± SD indicated.

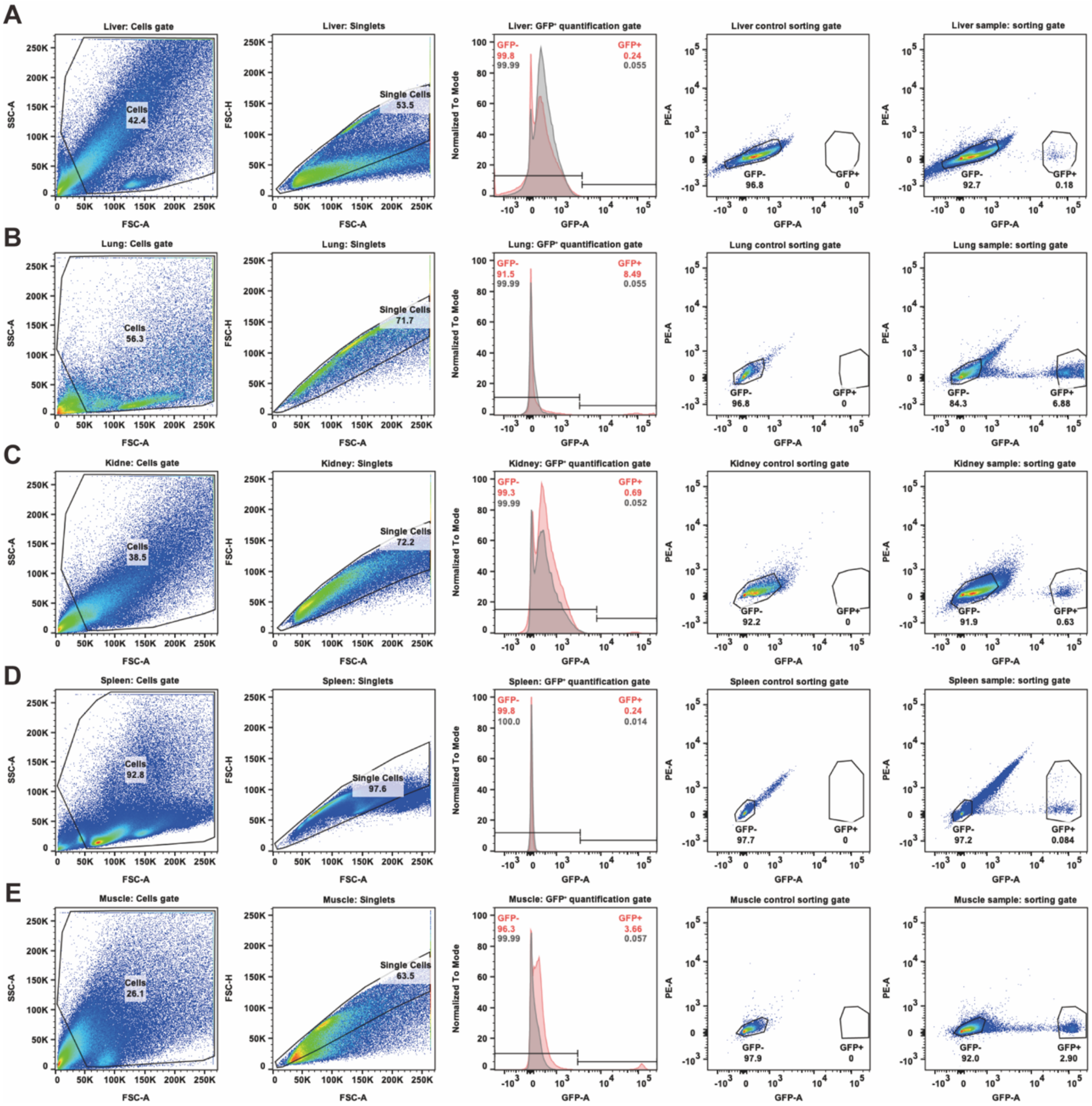

Supplementary Figure S2. Representative GFP gating across tissue-derived single-cell suspensions.

Representative GFP gating examples are shown for liver (A), lung (B), kidney (C), spleen (D) and muscle (E) single-cell suspensions from PDGFRα-eGFP mice and matched controls. For each tissue, pseudo-colour plots show GFP$^{-}$ and GFP$^{+}$ gates used for FACS sorting in control and PDGFRα-eGFP samples. Histogram plots show the GFP fluorescence threshold used to quantify GFP$^{+}$ cell frequency as a percentage of singlet events. Red histograms represent PDGFRα-eGFP tissue-derived samples and grey histograms represent matched controls.

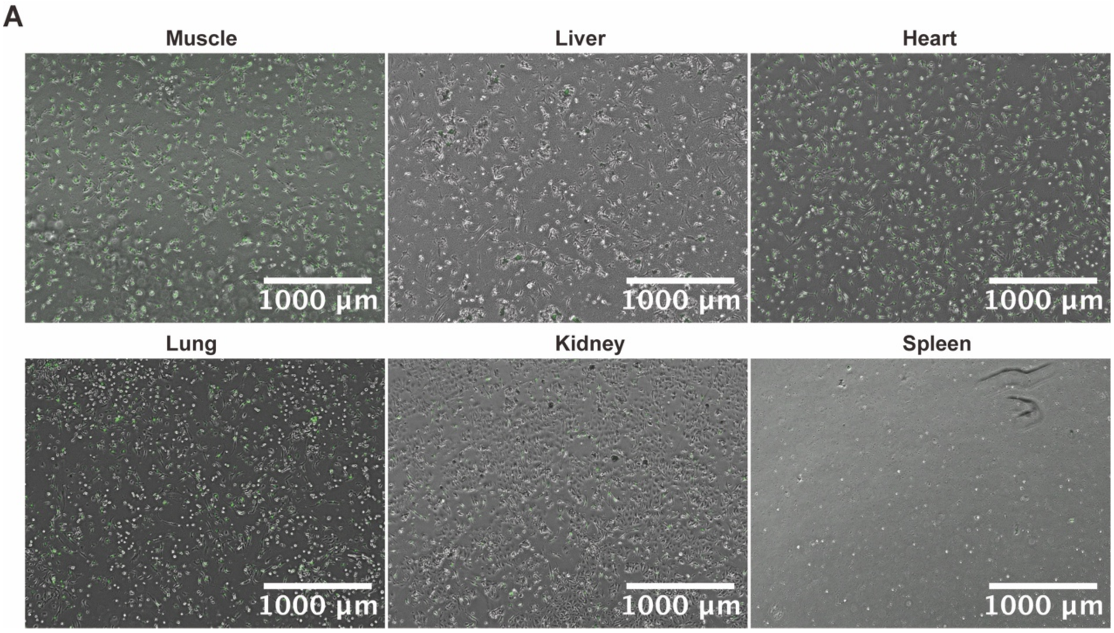


Supplementary Figure S3. GFP signal in short-term cultures derived from tissue single-cell suspensions of PDGFRα-eGFP mice.

Representative brightfield and GFP overlay images of short-term cultured single-cell suspensions derived from muscle, liver, heart, lung, kidney and spleen of PDGFRα-eGFP mice. GFP-positive cells are shown in green. Scale bars, 1000 µm.

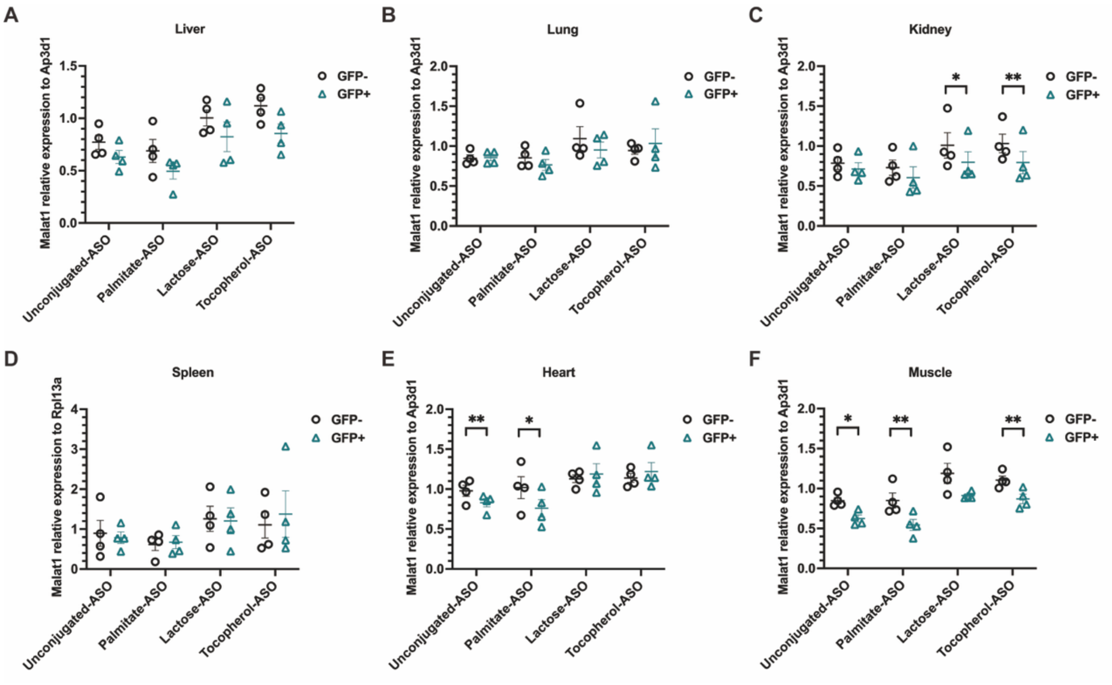


Supplementary Figure S4. Direct comparison of GFP⁺ and GFP⁻ pharmacodynamic responses following ASO treatment.

(A–F) Malat1 expression in GFP⁻ and GFP⁺ fractions sorted from liver (A), lung (B), kidney (C), spleen (D), heart (E) and muscle (F) after subcutaneous administration of unconjugated ASO, palmitate-ASO, lactose-ASO or tocopherol-ASO. Expression was normalised to *Ap3d1*, except for spleen samples, which were normalised to *Rpl13a*, and further normalised to the corresponding PBS-treated group within each sorted population. Each point represents an individual mouse; bars indicate mean ± SEM. Statistical comparisons between GFP⁺ and GFP⁻ populations within each treatment group were performed using paired two-tailed t-tests. *$P < 0.05$, **$P < 0.01$.

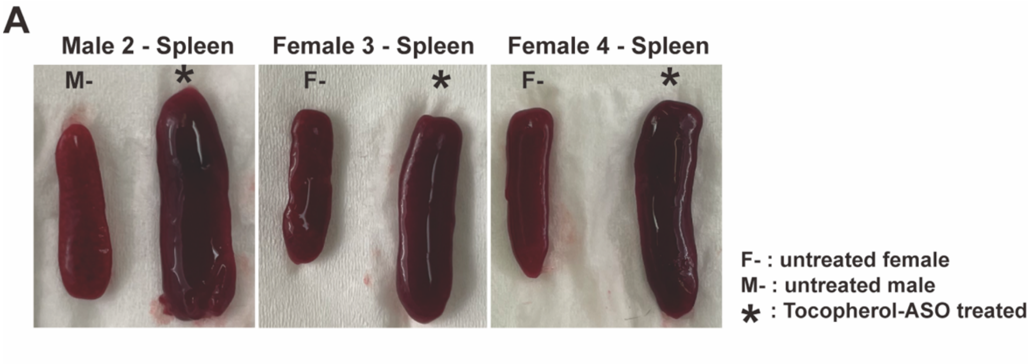


Supplementary Figure S5. Spleens from additional tocopherol-ASO-treated mice.
Gross spleen images from additional tocopherol-ASO-treated mice and age-matched untreated controls. One male and two female tocopherol-ASO-treated mice are shown with untreated controls of the corresponding sex.

**Supplementary Table 1:** ASO sequences and calculated and observed molecular masses. Observed molecular masses were determined by LC–MS following deconvolution of the mass spectra. Asterisks (*) indicate phosphorothioate (PS) internucleotide linkages; underlined nucleotides are 2′-*O*-methoxyethyl (2′-MOE)-modified, dN denotes a DNA nucleotide, and all cytosine residues are 5-methylcytosine (5mC).

| ASO Conjugate | Sequence (5'-3') | Calculated mass (Da) | Observed mass (Da) |
|---|---|---|---|
| none | A*T*G*5mC*A*dT*dT*d5mC*dT*dA*dA*dT*dA*dG*d5mC*A*G*5mC*A*G | 7227 | 7227 |
| Palmitoyl | A*T*G*5mC*A*dT*dT*d5mC*dT*dA*dA*dT*dA*dG*d5mC*A*G*5mC*A*G | 7644 | 7644 |
| Tocopheryl | A*T*G*5mC*A*dT*dT*d5mC*dT*dA*dA*dT*dA*dG*d5mC*A*G*5mC*A*G | 7925 | 7924 |
| Lactosyl | A*T*G*5mC*A*dT*dT*d5mC*dT*dA*dA*dT*dA*dG*d5mC*A*G*5mC*A*G | 7836 | 7838 |
